# Viral envelope proteins fused to multiple distinct fluorescent reporters to probe receptor binding

**DOI:** 10.1101/2023.10.23.563555

**Authors:** Ilhan Tomris, Roosmarijn van der Woude, Rebeca de Paiva Droes Rocha, Alba Torrents de la Peña, Andrew B. Ward, Robert P. de Vries

## Abstract

Enveloped viruses carry one or multiple proteins with receptor binding functionalities. Functional receptors can either be glycans, proteinaceous or both, recombinant protein approaches are instrumental to gain more insight into these binding properties. Visualizing and measuring receptor binding normally entails antibody detection or direct labelling, whereas direct fluorescent fusions are attractive tools in molecular biology. Here we report a suite of different fluorescent fusions, both N- and/or C-terminal, for influenza A virus hemagglutinins and SARS-CoV-2 spike RBD. The proteins contained a total of three or six fluorescent protein barrels and were applied directly to cells to determine receptor binding properties.

## Introduction

The surfaces of influenza A and SARS-CoV-2 viruses are decorated with glycoproteins that mediate receptor binding. For influenza A, hemagglutinin (HA), a homotrimer type 1 transmembrane glycoprotein, mediates receptor binding through interaction with α2-3 or α2-6-linked sialylated glycan structures on host cells [1-4]. Avian H3 and H5 viruses recognize only α2-3 linked sialic acid, preferably on branched N- and O-glycans [4-6]. Human H1 and H3 display specificity to Neu5Acα2-6Gal (α2-6 linked sialic acid), albeit with bias to certain features, such as contemporary H3 recognizing longer oligosaccharides with poly-LacNac repeats [6-10], whereas seasonal H1 shows preference to shorter structures [2, 5, 6, 11].

For SARS-CoV-2, the S spike glycoprotein mediates host cell receptor interaction using its receptor binding domain (RBD) [12, 13]. The S spike glycoprotein is composed of a S1 and S2 subunit, the RBD is a region located within the C-terminal domain (CTD) of the S1 subunit and directly interacts with the angiotensin-converting enzyme 2 (ACE2) receptor. Hereafter proteolytic cleavage by TMPRSS2 primes the S2 subunit, enabling membrane fusion and viral entry into host cells [14]. Other human coronaviruses also utilize the RBD located in the CTD for receptor binding, such as NL63, TGEV, PRCV, MERS and SARS [15]. Furthermore, many coronaviruses (TGEV, BCoV, MERS, HKU1 and OC43) possess an evolutionarily conserved glycan-binding region located in the N-terminal domain of the S1 subunit, similarly shown for SARS-CoV-2 [16-18]. For most influenza A and Coronavirus strains receptor binding specificities are not fully defined yet, therefore we need straightforward tools to study receptor binding properties.

The discovery of the naturally occurring green fluorescent protein (GFP), derived from Aequorea Victoria, has enabled the characterization, visualization, and protein localization within cellular processes [19-23]. The fusion of GFP to proteins of interest (P.O.I.) with molecular engineering enables the expression in a 1:1 ratio, thus allowing the use of quantitative approaches [24]. Multiple spectral variants have been developed, spanning a wide range on the visible spectrum (www.fpbase.org), allowing multiparameter measurements or advanced microscopy techniques [25-28]. Furthermore, the expression of recombinant fluorescent proteins may shed (more) information about the functionality of the protein, e.g, receptor-binding. We designed and expressed recombinant fluorescent HA and RBD proteins that allow for direct observation of receptor-binding interactions. The placement of a fluorescent protein (FPs) in relation to a P.O.I. may influence its functionality, e.g. protein folding and interference of target/binding domains [29]. Cytoplasmic proteins allow for FP fusion at the NH_2_- (N-terminal) or COOH-terminus (C-terminal), since proteins are able to fold with the N-terminal and C-terminal exposed on the surface of the protein, rather than being buried [30]. For transmembrane (glyco)proteins the placement of the FP can be restrictive towards expression and functionality [31]. HA proteins from influenza A viruses are such membrane-bound proteins, with the stem domain (C-terminus) facilitating the linkage towards the transmembrane domain. We have previously shown that the HA C-terminus tolerates the fusion of superfolder GFP (sfGFP) [11]. The N-terminus of HA folds downwards, next to the C-terminus, indicating a potential tolerance for a fluorescent fusion protein (FFP). Recombinant HAs were created with a GCN4 trimerization domain, Strep-Tag (TwinStrep, TS), with or without a sfGFP at the N- and/or C-terminus (Fig. 1). Additionally, we also generated recombinant HA with an improved palette of FPs (mTagBFP2, mTurquoise2, mOrange2, mCherry2, mPlum) with varying spectral properties (S1 Table), enabling multiparameter imaging/measurement. Furthermore, the genetic fusion of GFP appeared to play a role in increased expression yields [32, 33], therefore characterization of expression using different FPs is warranted.

**Figure 1.**
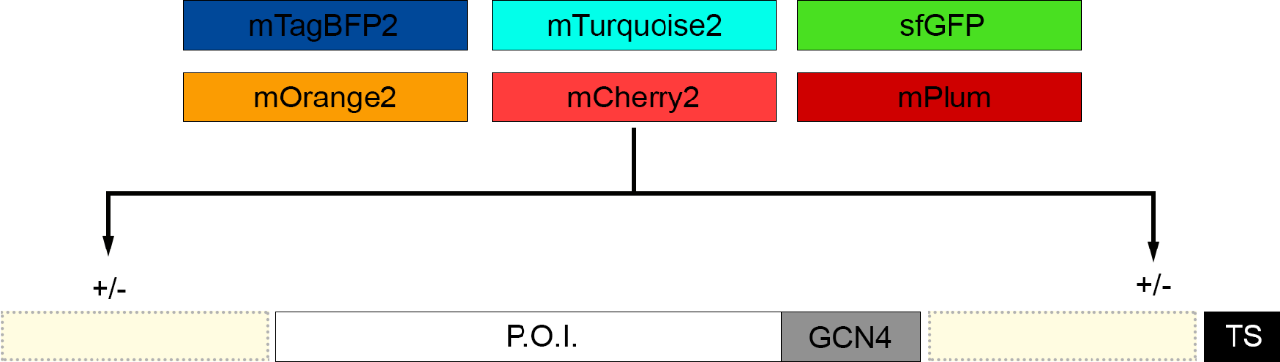
Designing fluorescent fusion proteins allows for visualization/detection with imaging techniques. The fluorescent probe sfGFP was cloned upstream (N-terminal) and/or downstream (C-terminal) of the protein of interest (P.O.I.). Fluorescent probes with different spectral properties were also cloned for utilization in multi-dye approaches.

## Results

### Genetic fusion of an improved palette of fluorescent influenza HA proteins

To demonstrate the tolerance of N-terminally linked FP we generated recombinant HAs in an expression plasmid containing A/Puerto Rico/8/1934 (A/PR/8/34) and A/Singapore/INFIMH-16-0019/2016 (A/SG/16). A/Puerto Rico/8/1934 is a model strain commonly used as reference, with A/SG/16 being difficult to express potentially related to prolonged Golgi retention [34]. We also created the N&C-terminal fusion and compared both with our previously made C-terminal FP construct [11]. Both the N- and N&C-terminal fusions were well tolerated as determined by Western blot and direct measurement of fluorescent in the cell culture supernatant (Fig. 2, S1 Fig. and S1 table). In certain instances, the N(&C)-terminal fusion enhanced the expression in comparison to the C-terminal variants (e.g. mCherry2-A/PR/8/34, Fig. 2). For mTagBFP2 expression was lower in general as observed with Western blot, even though fluorescence signal could be measured. Additionally, HA fusion-proteins with a FP at both ends (mTagBFP2, sfGFP, and mOrange2) appear to display the highest fluorescent capability, albeit not doubling the fluorescence intensity. Nonetheless, a fluorescent palette of recombinantly expressed influenza A hemagglutinin tolerates FFP at both termini, whilst also increasing protein expression yields for difficult to express protein (A/SG/16).

**Figure 2.**
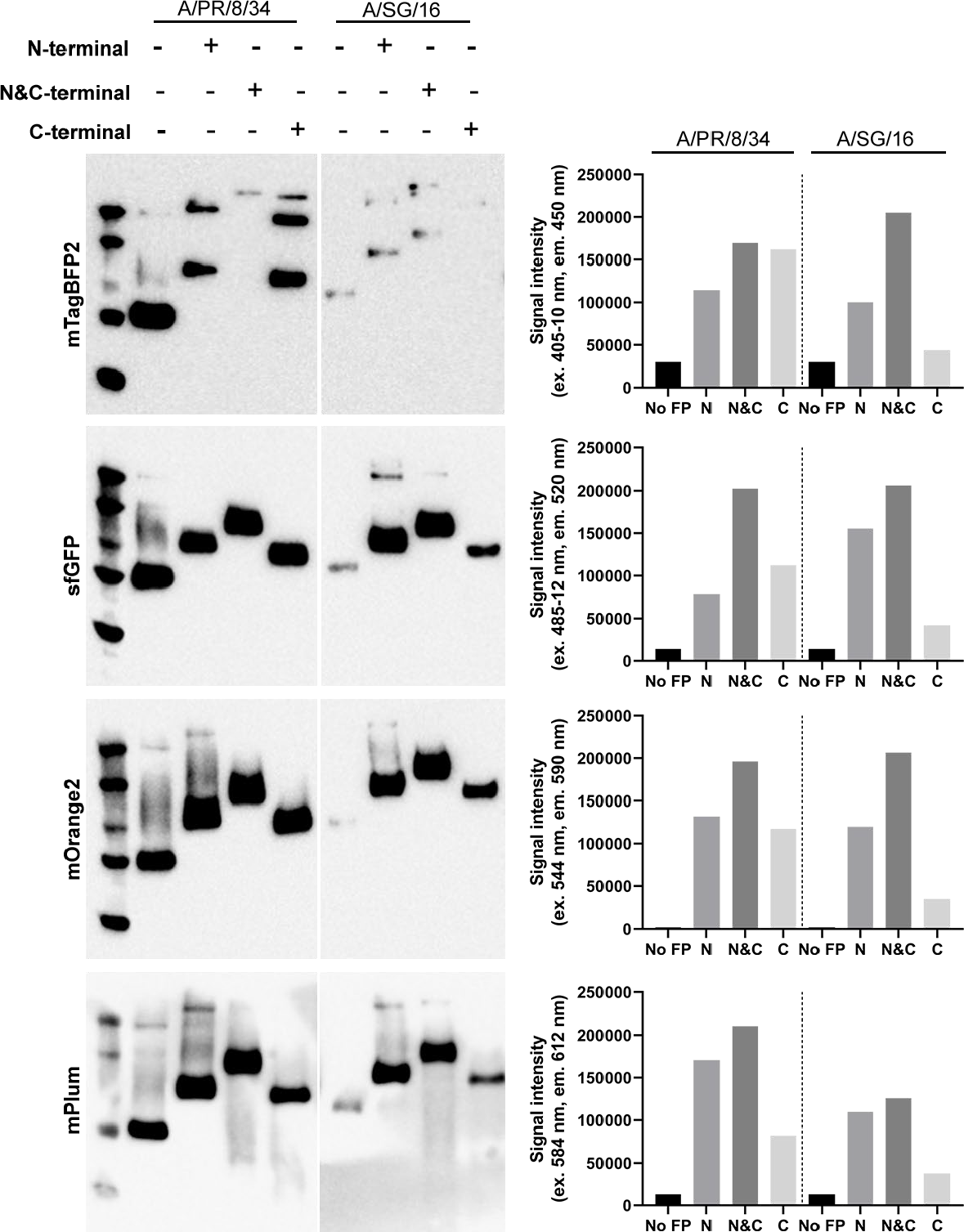
Palette of fluorescent HA expression. A/PR/8/34 and A/SG/16 expressed with different FFP (mTagBFP2, sfGFP, mOrange2, mPlum), whereby the FP was fused at the N-, N&C or C-terminus. Expression was characterized with Western-Blot and functionality of the fluorescent probe was verified with a fluorescence reader.

### Red fluorescent proteins for an improved fluorescent palette

Red-shifted FPs are desired for imaging applications, due to decreased light scattering in tissues and separation of autofluorescence [35]. We thus attempted the expression of several additional far-red fluorescent probes to improve our palette, however, these FPs completely abrogated protein expression or the FFP was no longer fluorescent (S2 table). Several FPs that have been reported as monomeric, appear to possess oligomeric tendencies (mNeptune, mCardinal and mKate2) [36], potentially related to the lack of success with our HA fusions. Monomerization of FPs is crucial, since oligomerization may have adverse effects, such as aggregation and transport resulting in containment within cellular compartments. Eventually, we managed to express mPlum-labelled HAs, which also increased the expression yield albeit possessing a lower brightness (Fig. 2 and S1 table)

### Folding properties of the P.O.I. are not negatively influenced by fluorescent fusion partners

A wide palette of FP fused to recombinantly expressed HA appears to be functional in terms of expression and fluorescence, however, these new fluorescent tools require antigenic and structural verification. To assess the antigenicity an ELISA assay was performed with conformation-dependent stalk binding antibodies. Furthermore, to expand on the applicability of the generated fluorescent tools, HA from additional influenza strains were included and the termini were fused to mOrange2, which was the best expressing and highly fluorescent fusion-protein. For A/PR/8/34 and A/Vietnam/1203/2004 (A/VN/1203/04) stalk antibody CR6261 was utilized [37], for A/SG/16, A/Hong Kong/8/1968 (A/HK/8/68) and A/duck/Ukraine/1963 (A/duck/UA/63) CR8020 [38]. ELISA data visualizes that for each HA strain fused to mOrange2, at the N-, N&C- and C-termini, no difference was observed, indicating that the folding properties of the P.O.I. are not influenced by the fusion-partner (Fig. 3A and S2. Fig). Subsequent characterization was performed with negative stain electron microscopy (ns-EM) to assess whether the proteins had a native-like structure and contained three or six barrels. When the sfGFP was present at the N- or C-terminus, three sfGFP barrels were observed and the protein was native-like (Fig. 3B and D). The sfGFP at the N-terminus folds downwards towards the C-terminal region, which directly faces the transmembrane domain (TMD) (Fig. 3D) [11]. For the N&C-terminal version six barrels were observed and the HA had a native-like structure (Fig. 3C), therefore, confirming the tolerance of an N-terminal FP when using a transmembrane-bound protein, without influencing the protein folding capacity. However, the potential interference of target/receptor binding for its biological functionality also requires assessment.

**Figure 3.**
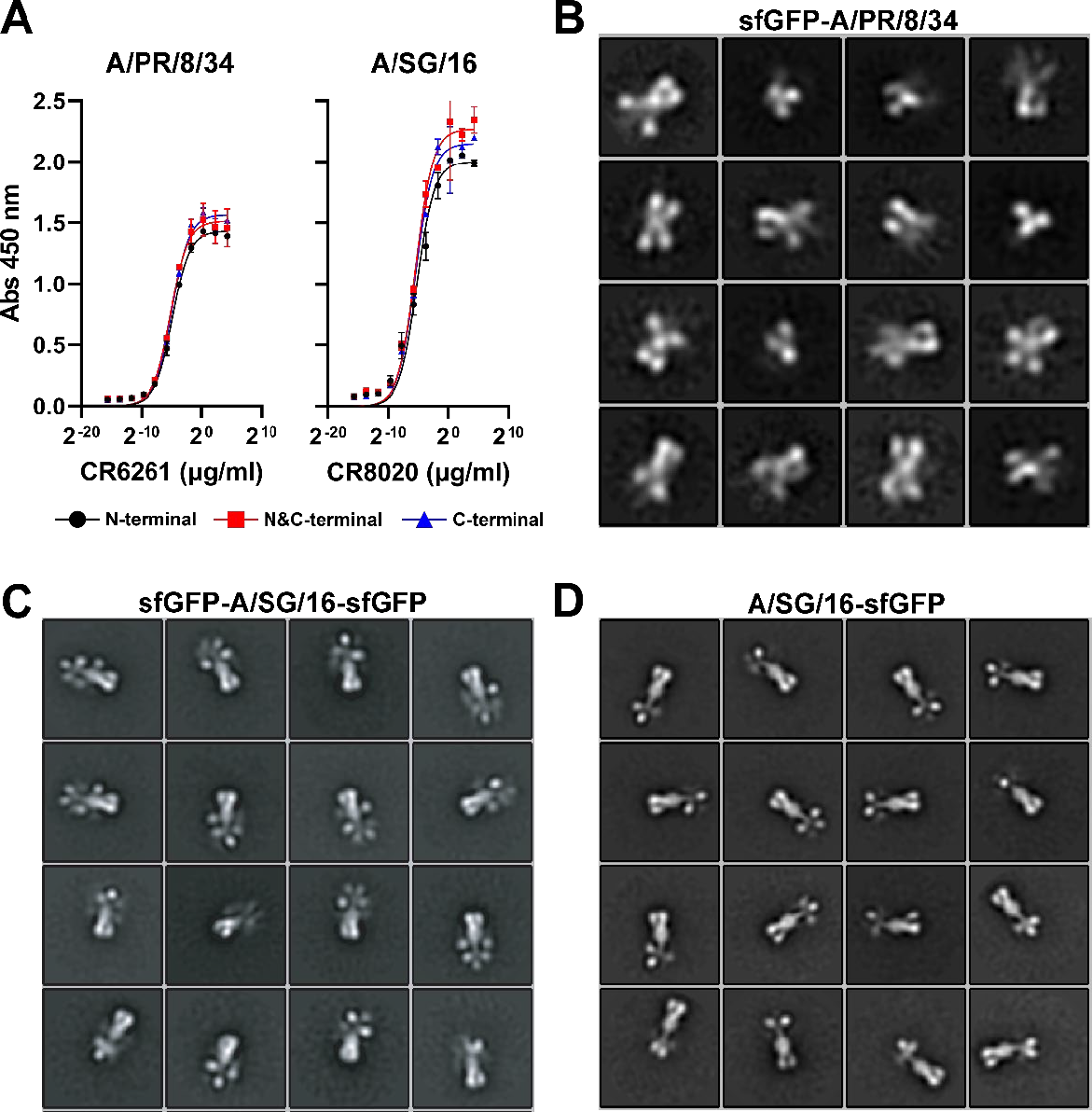
Fluorescent tools are antigenically and structurally similar. (A) ELISA assay to verify antigenicity of fluorescent fusion-protein (A/PR/8/34 and A/SG/16). Conformation-dependent stalk antibodies CR6261 and CR8020 utilized to verify conformation of recombinant HA in a concentration-dependent manner. (B) A/PR/8/34 with an N-terminal sfGFP folding downwards to the TMD. (C) A/SG/16 with an N&C-terminal sfGFP, six barrels visibly facing the C-terminus, therefore tolerating FPs at both ends and folding properly. (D) A/SG/16 with a C-terminal sfGFP in the tertiary conformation.

### Receptor binding functionality of hemagglutinin fluorescent fusion protein

Antigenicity and folding properties of the recombinantly expressed proteins are not influenced by an FFP. Therefore, to verify the biological functionality of these fluorescent recombinantly expressed HAs; further characterization with influenza-susceptible cell line Madin-Darby canine kidney (MDCK) [39] and Raji cells was performed. With fluorescence activated cell sorting (FACS) binding for A/PR/8/34, A/VN/1203/04, A/HK/1/68, A/SG/16 and A/duck/UA/63 was detected verifying the biological function of the protein, all proteins were expressed with a mOrange2 reporter (Fig. 4). For A/PR/8/34, A/VN/1203/04, A/duck/UA/63 and A/SG/16 the N&C-terminally labelled fusion proteins appear to have a higher signal compared to the N- or C-terminally labelled variants. The difference between N- and N&C-variants ranged from 1.18x to 1.87x, whilst between the C- and N&C-variants the difference ranged from 1.18x to 2.3x, potentially related to differences in protein maturation and folding of the mOrange2 within the individual HA trimers. Interestingly, A/HK/1/68 displayed different binding properties, dependent on the localization of the fusion protein, with the N-terminal FP being detrimental for receptor binding. For A/SG/16, fluorescence signal was higher in comparison to the MDCKs, which is potentially related to differences in glycan presentation on these cells [40-42].

**Figure 4.**
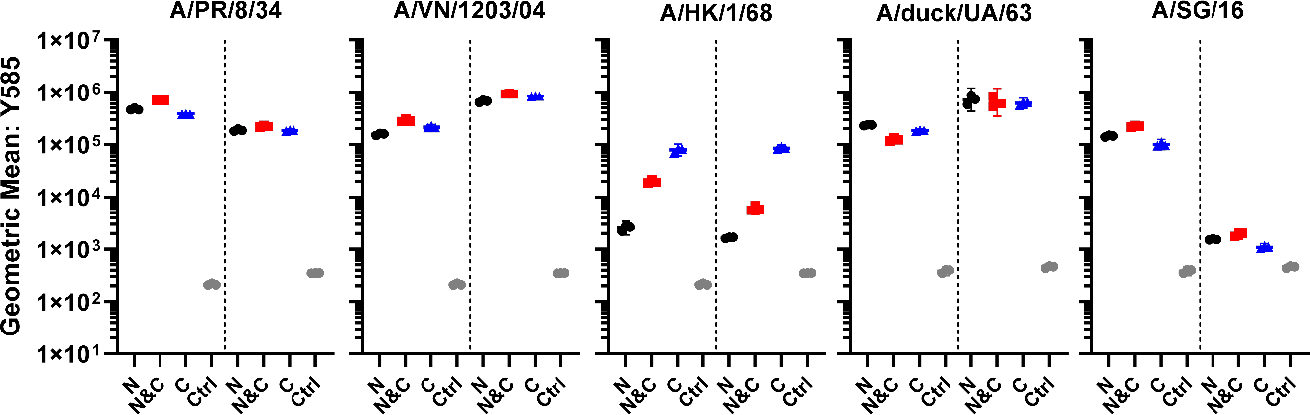
FFP receptor binding functionality. FACS performed with Rajis and MDCKs, in each graph Rajis are depicted on the left and MDCKs on the right. A/PR/8/34, A/VN/1203/4, A/SG/16, and A/duck/UA/63 display receptor binding with minor differences between the N-, N&C- and C-labelled variants. The presence of three additional mOrange2 barrels did not necessarily increase the fluorescence intensity two-fold. Ranging from 1.18x to 1.87x difference in fluorescence intensity between N- and N&C-variants and from 1.18x to 2.3x for the C- and N&C-variants. N-terminal mOrange2 displays detrimental binding for A/HK/1/68, whilst A/SG/16 signal is less in MDCK in relation to Rajis, possibly related to differences in glycan display. Control (Ctrl).

### Multiparameter characterization of fluorescent palette of hemagglutinin fusion protein

Multiparameter experiments were conducted with N&C-labelled FFP A/PR/8/34 (mTagBFP2), A/HK/1/68 (sfGFP), A/duck/UA/63 (mOrange2) and A/SG/16 (mPlum), to assess whether these fluorescent tools enable rapid characterization of binding. All FFPs from different strains were first measured individually to generate a compensation matrix for the spectral overlap of the FPs, followed by a simultaneous incubation using Rajis or MDCKs (Fig. 5). Separation of the individual channels, with HAs from different strains was possible, confirming the possibility of multiparameter measurement.

**Figure 5.**
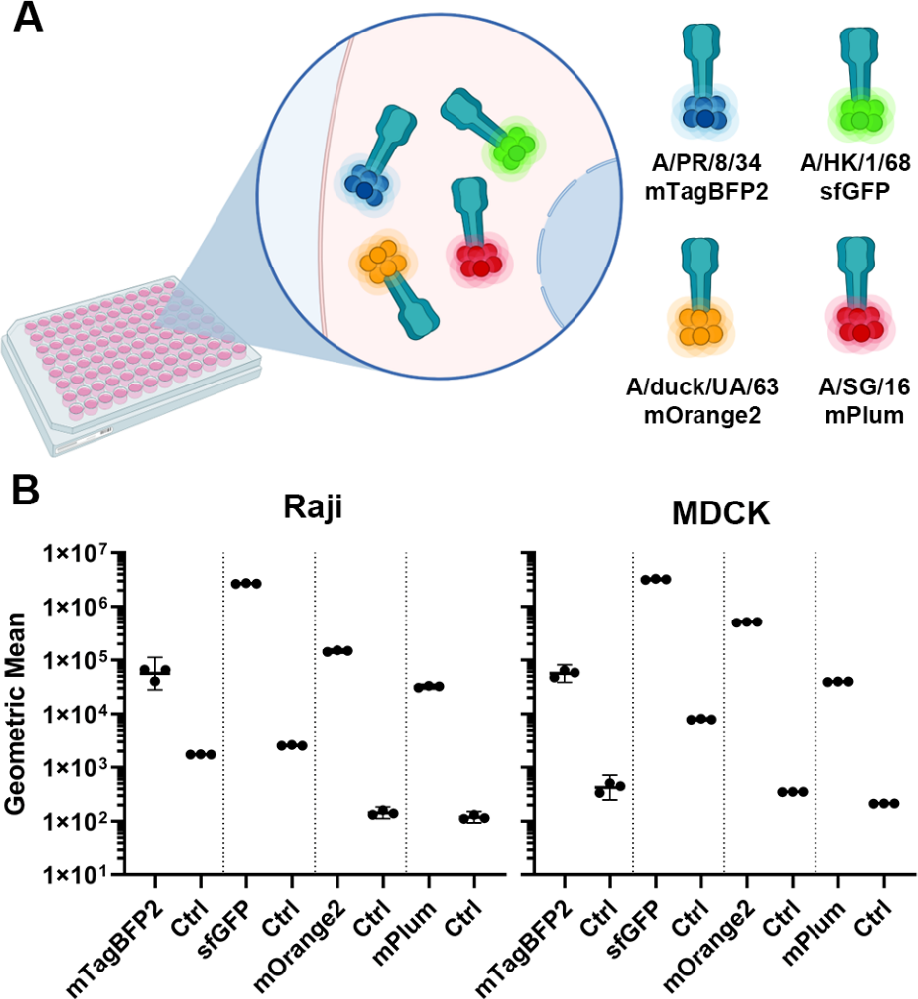
Multiparameter quantification. FACS performed with Rajis and MDCKs, in each graph Rajis are depicted on the left and MDCKs on the right. N&C-labelled FFP A/PR/8/34 (mTagBFP2), AHK/1/68 (sfGFP), A/duck/UA/63 (mOrange2) and A/SG/16 (mPlum) measurement displays the possibility for rapid measurement of different receptor-binding proteins on a certain substrate. Control (Ctrl).

### SARS-CoV-2 RBD fluorescent palette multiparameter characterization

To confirm whether our fluorescent tools were viable for other receptor binding proteins, unrelated to influenza, we replaced the HA open reading frame (ORF) with the ORF of SARS-CoV-2 RBD. Like HA, differences in expression were observed dependent on the FP (Fig. 6A). The N&C-terminal variant of mTagBFP2, mTq2, sfGFP mOrange2 and mPlum enhanced the expression in relation to the N- or C-labelled proteins. However, only for the N-terminal variant of mCherry2 expression was observed for SARS-CoV-2 RBD, but not when fused C-terminally. SARS-CoV-2 RBD expression appeared to decrease in general when fused to mTagBFP2 and mTq2, whilst expression of non-fluorescent SARS-CoV-2 RBD was higher. Incidentally, fusions with sfGFP, mOrange2 and mPlum did improve the expression and significant fluorescence was measured (Fig. 6B). Biological functionality of these FFPs were verified with Vero E6 cells, which is a relevant cell-line for SARS-CoV-2 [43]. The compensation matrix for HAs was utilized to correct for the spectral overlap. The SARS-CoV-2 RBDs labelled with different FPs appear to bind efficiently to the Vero E6 cells and it was possible to separate the individual channels. Similar to the multi-color HAs, SARS-CoV-2 RBDs displayed separation power of the differently labelled proteins. Therefore, demonstrating the functionality of these fluorescent tools for rapid multiparameter measurement.

**Figure 6.**
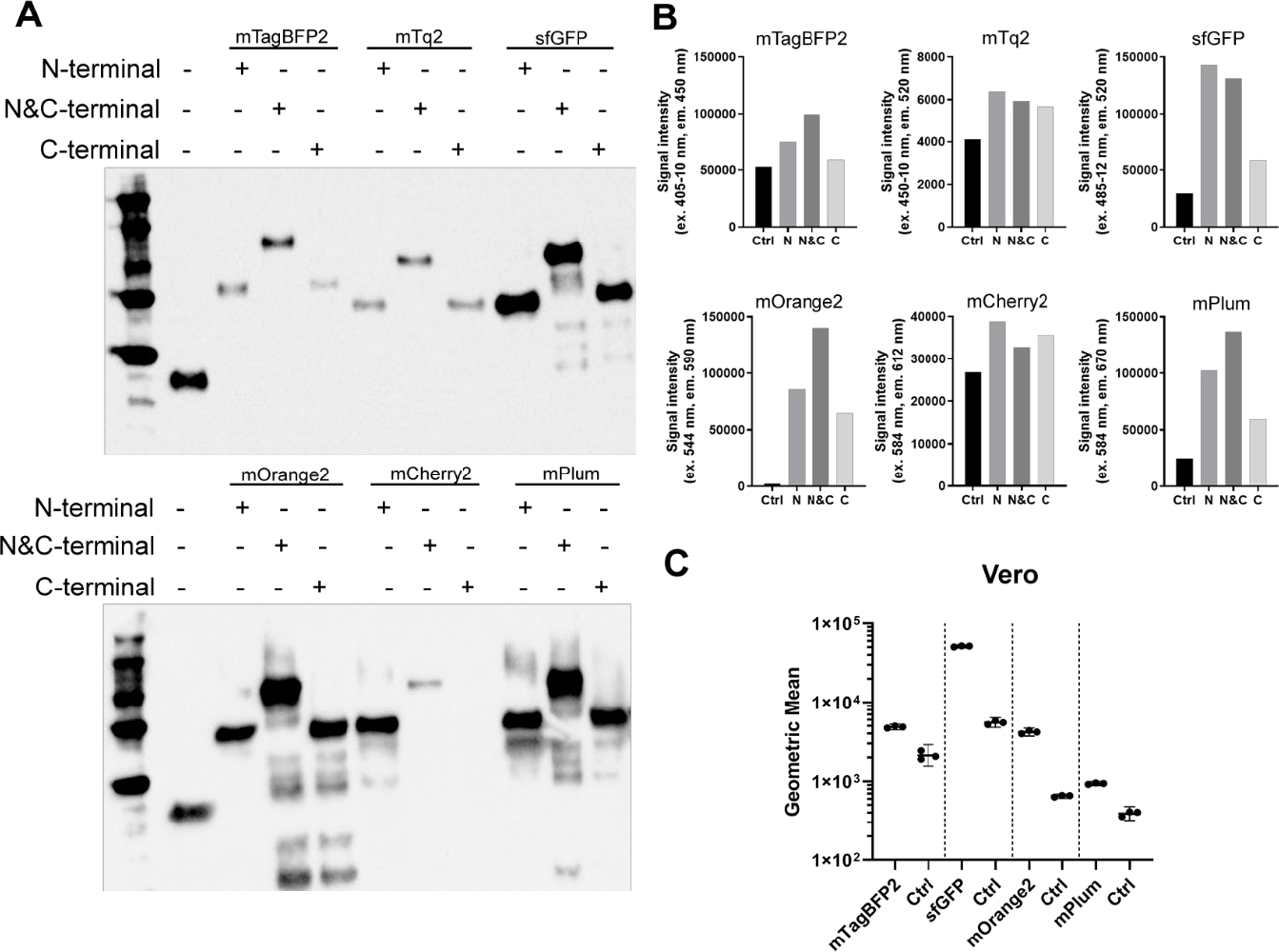
Fluorescent palette of SARS-CoV-2 RBD. (A) SARS-CoV-2 RBD fused to FPs (mTagBFP2, mTq2, sfGFP, mOrange2, mCherry2 and mPlum) N-, N&C- and C-terminally. The N&C labelled variants for all FPs, except mCherry2, resulted in the highest expression yield. (B) Fluorescence output was measured, the fluorescence signal intensities appeared to overlap with the expression profiles observed with Western blot, except for mTq2 and mCherry2. (C) N&C-labelled SARS-CoV-2 RBD FFP was utilized to assess biological functionality using Vero E6 cells.

## Discussion

Fluorescence techniques have become invaluable tools in the field of biotechnology. FPs have great advantages, in comparison to synthetic dyes as the genetic introduction enables imaging of live cells, organelles, single molecules and monitoring of dynamic interactions with other proteins or glycans [11, 44]. FPs have different advantages and limitations; thus, a proper selection is necessary to meet the experimental requirements. The selection of a FP is based on certain crucial factors, such as: spectral properties, quantum yield, brightness, maturation rate, fluorescence lifetime, monomeric character, and fidelity in fusions [19, 29, 45-47]. Here we created a set of fluorescent tools that span the visible spectrum using mTagBFP2, mTq2, sfGFP, mOrange2, mCherry2 and mPlum. The FPs mTagBFP2 and mTq2 are described as the brightest and most photostable blue/cyan dyes [48, 49], with sfGFP showing high brightness and folding efficiency [50]. In the orange-red spectrum, mOrange2 [51] has high photostability and mCherry2 [52] is a dye widely used in live-cell imaging for its high brightness and stability, monomeric nature and fast maturation. Certain fusion variants (A/PR/8/34, A/SG/16 and SARS-CoV-2 RBD) with mCherry2 displayed lower expression in comparison to mOrange2 and sfGFP fusions, potentially related to previous reports of protein aggregation [53]. Therefore, we expanded our palette with mPlum, with a similar excitation wavelength as mCherry2, however, with a longer emission wavelength for greater tissue penetration [54, 55].

FPs are oligomeric in nature, wild-type GFP from *A. Victoria* is part of a heterotetrameric complex [24], coral and anemone FPs are tetrameric [56]. Certain mutations have been introduced into these FPs to eliminate the oligomerization, which has been successful for jellyfish-derived FPs, but not completely for coral-derived FPs [36]. During expression of fluorescent proteins certain problems may occur, such as weak signal, suppression of fluorescence, protein aggregation, incorrect localization and non-functional FFP [46]. Oligomeric tendencies of FPs can be evaluated with an organized smooth endoplasmic reticulum (OSER) assay, whereby oligomerization leads to reconfiguration of the ER [46, 57]. Interestingly, mTagBFP2 and superfolder GFP also display an oligomeric nature, however, we could express and utilize these proteins in our experiments with ease. Additionally, the expression of most RFPs was unsuccessful, however, we managed to obtain a functional probe, albeit with a low brightness. FP performance in fusions is dependent on more aspects than oligomerization, potentially influenced by each individual FP, cell type, mRNA stability, translation efficiency, stability of the FP or maturation-dependent hydrogen peroxide production [46, 52, 58].

Recombinant protein expression is a staple of molecular biology and the expression yield determines whether a P.O.I. is suitable for experimental purposes [33, 59]. FPs could thus enhance protein expression yields to characterize the functionality of proteins, whilst serving as an intrinsic fluorescent handle. All in all, here we display a set of fluorescent tools that span the visible spectrum. We emphasize that the FP and the location of the FP in relation to the P.O.I. can influence the expression yield, without hampering folding and receptor binding capacity. Thus, underlining the necessity of properly evaluating current expression systems and how expression can be enhanced with these probes. Additionally, we demonstrate the functionality of these fluorescent tools for rapid multiparameter measurement. In addition, these tools are also useful for competition assays with different fluorescent handles and enable identification of specific cell types within a mixed population when using proteins with different specificity [28].

## Material and Methods

### HA expression plasmid generation

Codon-optimized HA encoding sequences of A/Puerto Rico/8/34, A/Vietnam/1203/04, A/Hong Kong/1/1968, A/duck/Ukraine/1963 and A/Singapore/INFIMH-16-0019/2016 were cloned into the pCD5 expression as described previously [60]. Full length SARS-CoV-2 S spike 2 (GenBank: MN908947.3) encoding open reading frames (A kind gift of Rogier Sanders, Amsterdam Medical Centre, The Netherlands), the RBD subunit (SARS-2 319–541) was amplified using PCR as described previously [61]. Expression vector pCD5 was adapted, for N- and N&C-terminal vectors, the signal sequence was followed by a fluorescent protein open reading frame, the HA-encoding cDNA, GCN4-pII trimerization motif (KQIEDKIEEIESKQKKIENEIARIKK, a TEV cleavage site, a fluorescent reporter (N&C vectors) and a Strep-tag II (WSHPQFEKGGGSGGGSWSHPQFEK); IBA, Germany). The C-terminal vectors were adapted as described previously [11]. The fluorescent reporter open-reading frames (S3 table) were ordered from Addgene and cloned into the accepting vectors utilizing PCR and Gibson assembly. The mTurquoise2 open reading frame was provided by Joachim Goedhart.

## Protein Expression and purification

pCD5-HA- +/- GCN4-Fluorescent probe expression vectors were transfected into HEK293S GNT1(-) cells (which are modified HEK293S cells lacking glucosaminyltransferase I activity (ATCC® CRL-3022™)) with polyethyleneimine I (PEI) in a 1:8 ratio (μg DNA:μg PEI) as previously described [60]. The transfection mix was replaced after 6 hours by 293 SFM II suspension medium (Invitrogen, 11686029, supplemented with glucose 2.0 gram/L, sodium bicarbonate 3.6 gram/L, primatone 3.0 gram/L (Kerry), 1% glutaMAX (Gibco), 1.5% DMSO and 2mM valproic acid). Culture supernatants were harvested 5 days post-transfection. The HA expression was analysed with SDS-PAGE followed by Western-blot on PVDF membrane (Biorad) using α-strep-tag mouse antibodies 1:3000 (IBA Life Sciences). Additionally, fluorescence intensities were measured using a filter based PolarStar Omega plate reader. To detect the fluorescence signal, respective fluorescence wavelengths for excitation and emission were utilized (S4 table). Subsequently, HA proteins were purified with Sepharose Strep-Tactin beads (IBA Life Sciences) as previously described [60].

## Antigenicity of HA proteins

To assess the antigenicity of the HA proteins, 5 μg/ml of HAs with a mOrange2 fluorescent reporter was coated on MaxiSorp 96-wells plates overnight at 4°C. Plates were blocked for three hours with 3% BSA in PBS 0.1% Tween20. CR6261 and CR8020 antibodies were serially diluted 1:1 with a starting concentration of 20 μg/ml, followed by an incubation of one hour at room temperature. A secondary antibody goat-anti-human HRP (31410, Thermo Scientific) at 1:2.000 dilution was incubated at room temperature for one hour to detect the primary antibody. TMB substrate solution (34028, Thermo Scientific) was utilized to develop the plates and the reaction was stopped after 5 minutes using 2.5 M H_2_SO_4_. The absorbance was measured at 450 nm with a POLARstar Omega plate reader.

## Negative stain electron microscopy structural analysis

HA proteins in 100mM Tris, 150mM NaCl at 4°C were diluted to 0.025 mg/ml, deposited on 400 mesh copper negative stain grids for 10 s and doubled stained with 2% uranyl formate for 10 s and 30 s. The grids were imaged on a 120KeV Tecnai Spirit or 200KeV Tecnai T20 electron microscope with a LaB6 filament and a 4k x 4k TemCam F416 camera. Micrographs were collected using Legion [62] and then uploaded to Appion [63]. Particles were picked using DoGPicker [64] and further 2D classification was performed using Relion 3.0 [65].

## Biological activity and multiparameter flow cytometry

A/PR/8/34, A/VN/1203/04, A/HK/1/1968, A/duck/UA/1963 and A/SG/2016 were utilized in the flow cytometry measurements, whereby only the mOrange2 labelled proteins were used. Fluorescent HA proteins (100 μg/ml) were precomplexed with a primary α-strep-tag mouse antibody (50 μg/ml, 2-1507-001, IBA Life Sciences), a secondary rabbit-anti-mouse-HRP (25 μg/ml, NB7544, Novus Biologicals) and incubated for one hour at 4°C. For each single measurement precomplexed proteins were added to either 50.000 Raji or MDCK cells, followed by a one-hour incubation period at 4°C. Following the staining cells were washed with in FACS buffer (PBS, 0.5% BSA and 2 mM EDTA) and spin down at 200 rcf for 5 mins. Viability staining was performed with ViaKrome 808 viability dye (C36628, Beckman Coulter) 1:10.000 diluted in FACS buffer for 5 mins at 4°C, followed by centrifugation at 200 rcf for 5 mins. Flow cytometry experiments were performed on a Cytoflex LX and fluorescent signal was detected in the Y585 channel. Gating strategies for the cell population, singlets, time and viable cells were employed (S3 Fig.) and the fluorescent signal intensities were quantified in the Y585 channel and plotted in GraphPad v9, Geometric mean values with a 95% confidence interval. For the multiparameter HA measurements only N&C-terminally labelled FFP were utilized, A/PR/8/34 (mTagBFP2), A/HK/1/1968 (sfGFP), A/duck/UA/1963 (mOrange2) and A/SG/2016 (mPlum), proteins (100 μg/ml) were individually precomplexed with a primary α-strep-tag mouse antibody (50 μg/ml, IBA Life Sciences), a secondary rabbit-anti-mouse-HRP (25 μg/ml) and incubated for one hour at 4°C. Hereafter the HAs with different fluorescent reporters were first used for single stains for the compensation matrix of the flow cytometer, followed by a simultaneous stain with all HAs with different reporters. Fluorescence signal was measured in the V450, B525, Y585 and Y675 channel of the CytoFlex flow cytometer. Gating strategies for the cell population, singlets, time and viable cells were employed and the fluorescent signal intensities were plotted in GraphPad v9. Experiments with SARS-CoV-2 RBD (100 μg/ml) were performed similarly as the HAs, however, without any prior precomplexation. Additionally, staining was performed with Vero E6 cells.

## Acknowledgements

R.P.dV is a recipient of an ERC Starting Grant from the European Commission (802780). We thank the Netherlands Organization for Scientific Research (NWO) for the Rubicon Grant 45219118 to A.T.d.l.P.

## Notes

### Competing Interest Statement

The authors have declared no competing interest.

