## Supplementary figures for "Viral envelope proteins fused to multiple distinct fluorescent reporters to probe receptor binding"

**S1 Fig**

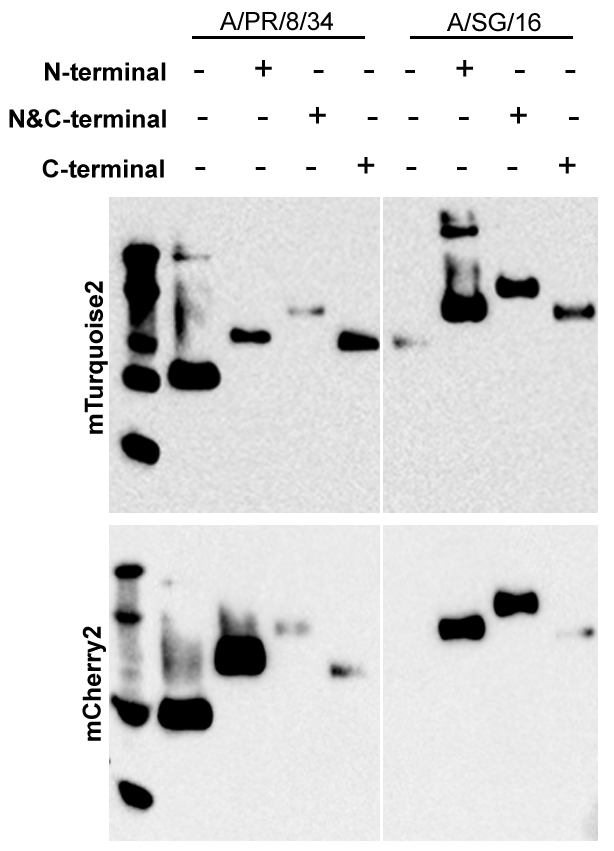

**S1 Fig. Fluorescent HA expression.** A/PR/8/34 and A/SG/16 expressed with mTq2 and mCherry2, whereby the FP was fused at the N-, N&C or C-terminus. Expression was characterized with Western-Blot.

**S2 Fig**

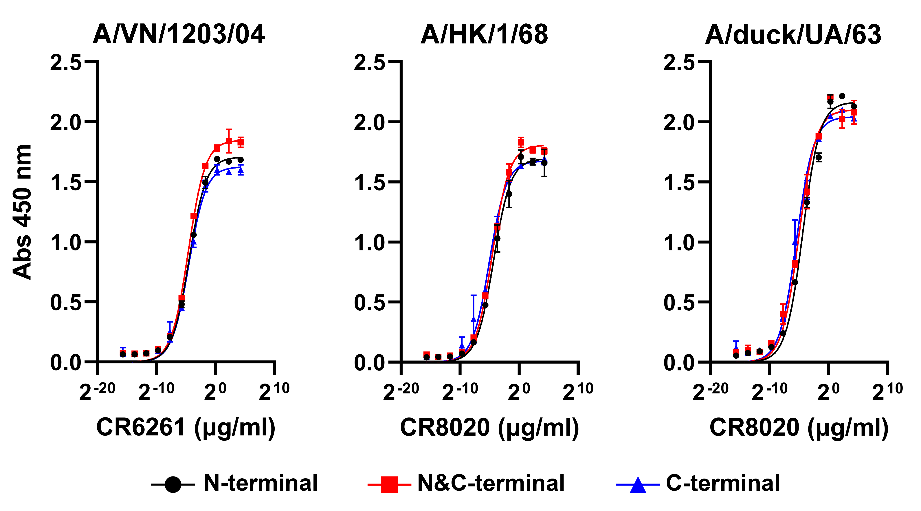

**S2 Fig. Antigenicity of FFPs.** HA from different strains utilized, A/VN/1203/4, A/HK/8/68 and A/duck/UA/63 with conformation-dependent antibody CR6261 or CR8020. Location of the fusion-partner did not influence folding characteristics.

**S3 Fig**

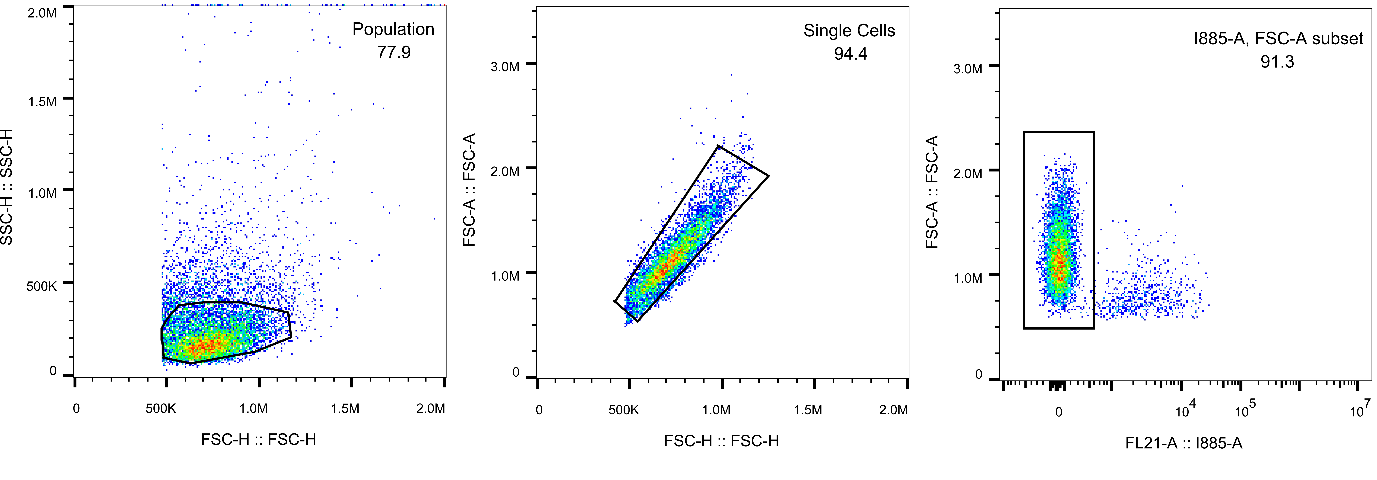

**S3 Fig. Gating strategies for flow cytometry.** Gating was performed to select for cell population, followed by singlets and then live-cells using ViaKrome 808 viability dye.

**S1 Table: Spectral properties of fluorescent proteins**

| **FP** | **Ex λ** | **Em λ** | **EC (M-1 cm-1)** | **QY** | **Brightness** | **Derived from** |
| --- | --- | --- | --- | --- | --- | --- |
| mTagBFP2 | 399 | 454 | 50,600 | 0.64 | 32.38 | Entacmaea quadricolor |
| mTurquoise2 | 434 | 474 | 30,000 | 0.93 | 27.9 | Aequorea victoria |
| sfGFP | 485 | 510 | 83,300 | 0.65 | 54.15 | Aequorea victoria |
| mOrange2 | 549 | 565 | 58,000 | 0.6 | 34.8 | Discosoma sp |
| mCherry2 | 589 | 610 | 79,400 | 0.22 | 17.47 | Discosoma sp |
| mPlum | 590 | 649 | 41,000 | 0.1 | 4.1 | Discosoma sp |

**S2 Table: Spectral properties of far-red fluorescent proteins**

| **FP** | **Ex λ** | **Em λ** | **Brightness** | **Derived from** | **Expression** | **Fluorescence** | **Ref.** |
| --- | --- | --- | --- | --- | --- | --- | --- |
| LSSmKate2 | 460 | 605 | 4.42 | Entacmaea quadricolor | No | x | [66] |
| FusionRed | 580 | 608 | 17.95 | Entacmaea quadricolor | yes | yes | [67] |
| mKelly2 | 598 | 649 | 7.74 | Entacmaea quadricolor | No | x | [36] |
| mGarnet | 598 | 670 | 8.55 | Entacmaea quadricolor | No | x | [68] |
| mNeptune | 600 | 650 | 13.4 | Entacmaea quadricolor | No | x | [69] |
| mCarmine | 603 | 675 | 5.81 | Entacmaea quadricolor | No | x | [70] |
| mCardinal | 604 | 659 | 16.53 | Entacmaea quadricolor | No | x | [71] |
| mMaroon | 609 | 657 | 8.8 | Entacmaea quadricolor | No | x | [72] |
| miRFP670 | 642 | 670 | 12.24 | Rhodopseudomonas palustris | No | x | [73] |
| miRFP670nano | 645 | 670 | 10.26 | Nostoc punctiforme | Yes | no | [74] |
| mIFP | 683 | 704 | 6.56 | Bradyrhizobium sp | No | x | [75] |

**S3 table: Open-reading frames fluorescent reporters**

| **Plasmid #** | **Fluorescent reporter** | **Plasmid #** | **Fluorescent reporter** |
| --- | --- | --- | --- |
| 57969 | mOrange2-Mito-7 | 54572 | mTagBFP2-pBAD |
| 56217 | mIFP-CytERM-N-17 | 54714 | mNeptune-pBAD |
| 54563 | mCherry2-C1 | 31909 | pBAD/HisD-LSSmKate2 |
| 54778 | FusionRed-Lifeact-7 | 104307 | LifeAct-mGarnet |
| 79987 | pmiRFP670-N1 | 54590 | mCardinal-N1 |
| 54554 | mMaroon-N1 | 119783 | pET-mKelly2 |
| 109486 | pcDNA3-mCarmine | 55966 | mPlum-ER-3 |
| 127427 | pBAD/His-miRFP670nano |  |  |

**S4 table: excitation and emission wavelength (nm) for fluorescence measurement**

| **Fluorescent reporter** | **Excitation (nm)** | **Emission (nm)** |
| --- | --- | --- |
| mTagBFP2 | 405-10 | 450 |
| mTurquoise2 | 450-10 | 520 |
| sfGFP | 485-12 | 520 |
| mOrange2 | 544 | 590 |
| mCherry2 | 584 | 612 |
| mPlum | 584 | 670 |
